# The electrically conductive pili of *Geobacter soli*

**DOI:** 10.1101/2020.01.09.901157

**Authors:** Shiyan Zhuo, Guiqin Yang, Li Zhuang

## Abstract

Electrically conductive pili (e-pili) enable electron transport over multiple cell lengths to extracellular environments and play an important role in extracellular electron transfer (EET) of *Geobacter* species. To date, the studies of e-pili have mainly focused on *Geobacter sulfurreducens* and the closely related *Geobacter metallireducens* because of their developed genetic manipulation systems. We investigated the role of *G. soli* pili in EET by directly deleting the pilin gene, *pilA*, which is predicted to encode e-pili. Deletion of *pilA*, prevented the production of pili, resulting in poor Fe(III) oxide reduction and low current production, implying that *G. soli* pili is required for EET. To further evaluate the conductivity of *G. soli* pili compared with *G. sulfurreducens* pili, the *pilA* of *G. soli* was heterologously expressed in *G. sulfurreducens*, yielding the *G. sulfurreducens* strain GSP. This strain produced abundant pili with similar conductivity to the control strain that expressed native *G. sulfurreducens* pili, consistent with *G. soli* as determined by direct measurement, which suggested that *G. soli* pili is electrically conductive. Surprisingly, strain GSP was deficient in Fe(III) oxide reduction and current production due to the impaired content of outer-surface *c*-type cytochromes. These results demonstrated that heterologous pili of *G. sulfurreducens* severely reduces the content of outer-surface *c*-type cytochromes and consequently eliminates the capacity for EET, which strongly suggests an attention should be paid to the content of *c*-type cytochromes when employing *G. sulfurreducens* to heterologously express pili from other microorganisms.

**IMPORTANCE:** The studies of electrically conductive pili (e-pili) of *Geobacter* species are of interest because of its application prospects in electronic materials. e-Pili are considered a substitution for electronic materials due to its renewability, biodegradability and robustness. Continued exploration of additional e-pili of *Geobacter soli* will improve the understanding of their biological role in extracellular electron transfer and expand the range of available electronic materials. Heterologously expressing the pilin genes from phylogenetically diverse microorganisms has been proposed as an emerging approach to screen potential e-pili according to high current densities. However, our results indicated that a *Geobacter sulfurreducens* strain heterologously expressing a pilin gene produced low current densities that resulted from a lack of content of *c*-type cytochromes, which were likely to possess e-pili. These results provide referential significance to yield e-pili from diverse microorganisms.

## Introduction

*Geobacter* species have a unique ecological niche due to their capability of extracellular electron transfer in diverse anaerobic environments (1). Electrically conductive pili (e-pili) are capable of transporting electrons along its length to extracellular environments and play an important role in the extracellular electron transfer of *Geobacter* species (2). e-Pili are required for Fe(III) oxides reduction by *Geobacter* species in sediments and soils because it can permit cells to make direct connections with insoluble electron acceptors multiple cell lengths from the cell surface (3, 4). e-Pili networks confer the ability of *Geobacter* species to produce high current densities and form thick biofilms in bioelectrochemical systems (1, 5). e-Pili of *Geobacter* species that function as electron donating partners can promote direct interspecies electron transfer (DIET) with methanogens to produce methane in anaerobic digestion (6, 7). In addition, e-pili serve as a sustainable electronic material with diverse potential applications (2, 8).

To date, most studies of e-pili have mainly focused on *Geobacter sulfurreducens* and the closely related *Geobacter metallireducens* because of their developed genetic manipulation systems (4, 9–11). The functions of pili and other proteins involved in extracellular electron transfer have been characterized in both *Geobacter* strains by evaluating the phenotype of gene deletions (4, 9, 12–15). However, the roles of pili of other microorganisms that have similar pilin monomer genes as *G. sulfurreducens* (16), especially difficult-to-culture microorganisms, are very difficult to study because of the lack of tools for genetic manipulation. Additionally, the conductivity of pili in these microorganisms is technically challenging to evaluate because of the tiny diameter of e-pili (ca. 3 nm). Another option to rapidly screen potential e-pili from diverse microorganisms is by heterologously expressing pilin genes in *G. sulfurreducens* in place of its native pilin gene (2, 16). Only If the heterologous pilin gene of *G. sulfurreducens* yields e-pili, the host will have the ability to generate high current densities on graphite electrodes (2, 16). In addition, the heterologous expression of pili from other microorganisms in *G. sulfurreducens* provides an alternative strategy to evaluate the conductivity of their pili in a common host (16, 17). For example, heterologously expressing the pilin gene of *G. metallireducens* in *G. sulfurreducens* yielded e-pili with 5,000-fold higher conductivity than its host (17).

*Geobacter soli*, distinct from *G. sulfurreducens* (18) and *G. anodireducens* (19), was the first *Geobacter* species isolated from forest soil (20). The physiological properties of *G. soli* are different from those of *G. sulfurreducens*, and these properties determine its special environmental significance, although both strains are highly similar based on phylogenetic analysis (20). For example, *G. soli* can anaerobically oxidize aromatic compounds, including phenol, benzoate and benzaldehyde, while *G. sulfurreducens* cannot (20, 21). In addition, *G. soli* has faster Fe(III) oxide reduction and higher current production than *G. sulfurreducens* (Fig. S1), which implies its stronger capacity of extracellular electron transfer compared to *G. sulfurreducens*. *G. soli* pili is truncated, similar to *G. sulfurreducens*, which is a key feature for microorganisms to assemble into e-pili (11, 22). The pilin gene of *G. soli* was predicted to encode e-pili by genome data analysis and phylogenetic analysis (22). Therefore, it is of interest to elucidate the role of *G. soli* pili in extracellular electron transfer, which might account for its special physiological characteristics.

In this work, the pilin gene of *G. soli* was directly deleted and heterologously expressed (i) to explore the role of *G. soli* pili in Fe(III) reduction and current production, (ii) to investigate the conductivity of *G. soli* pili and (iii) to further evaluate the conductivity of *G. soli* pili compared with *G. sulfurreducens* pili. Establishing an additional model of e-pili in *G. soli* will improve our understanding of their biological role in extracellular electron transfer and provide alternative electronic materials in biological applications.

## Results

### Pili of *G. soli* is electrically conductive

The pilin of *G. soli* has 95% amino acid sequence similarity and slightly higher aromatic amino acid content (10.8%) than *G. sulfurreducens* (9.8%) because it has an additional tyrosine at position 63 (Fig. 1). The conductivity of *G. soli* pili was directly measured by CP-AFM, and the results showed that it was half that of *G. sulfurreducens* (Fig. 2a, 2b, Fig. 5 and Fig. S2). The conductivity of *Geobacter* pili spans a very wide range, from 5,000-fold higher (*G. metallireducens*) to 150-fold lower (*G. uraniireducens*) compared with *G. sulfurreducens* (17, 23). Thus, we believed that the conductivity of *G. soli* pili was very comparable to that of *G. sulfurreducens* pili. Additionally, the diameter of *G. soli* pili was approximately 3 nm, the same as that of *G. sulfurreducens* pili (Fig. 2c).

**Fig. 1.**
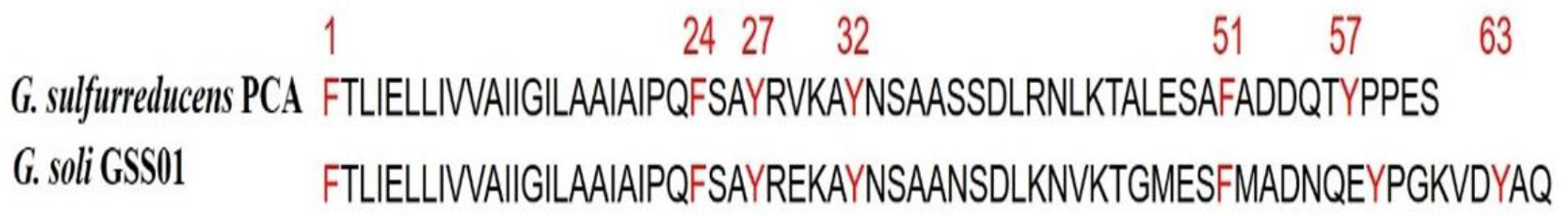
Alignment of PilA amino acid sequences of *Geobacter sulfurreducens* and *Geobacter soli*. The aromatic amino acids (F, phenylalanine; Y, tyrosine) are marked in red color.

**Fig. 2.**
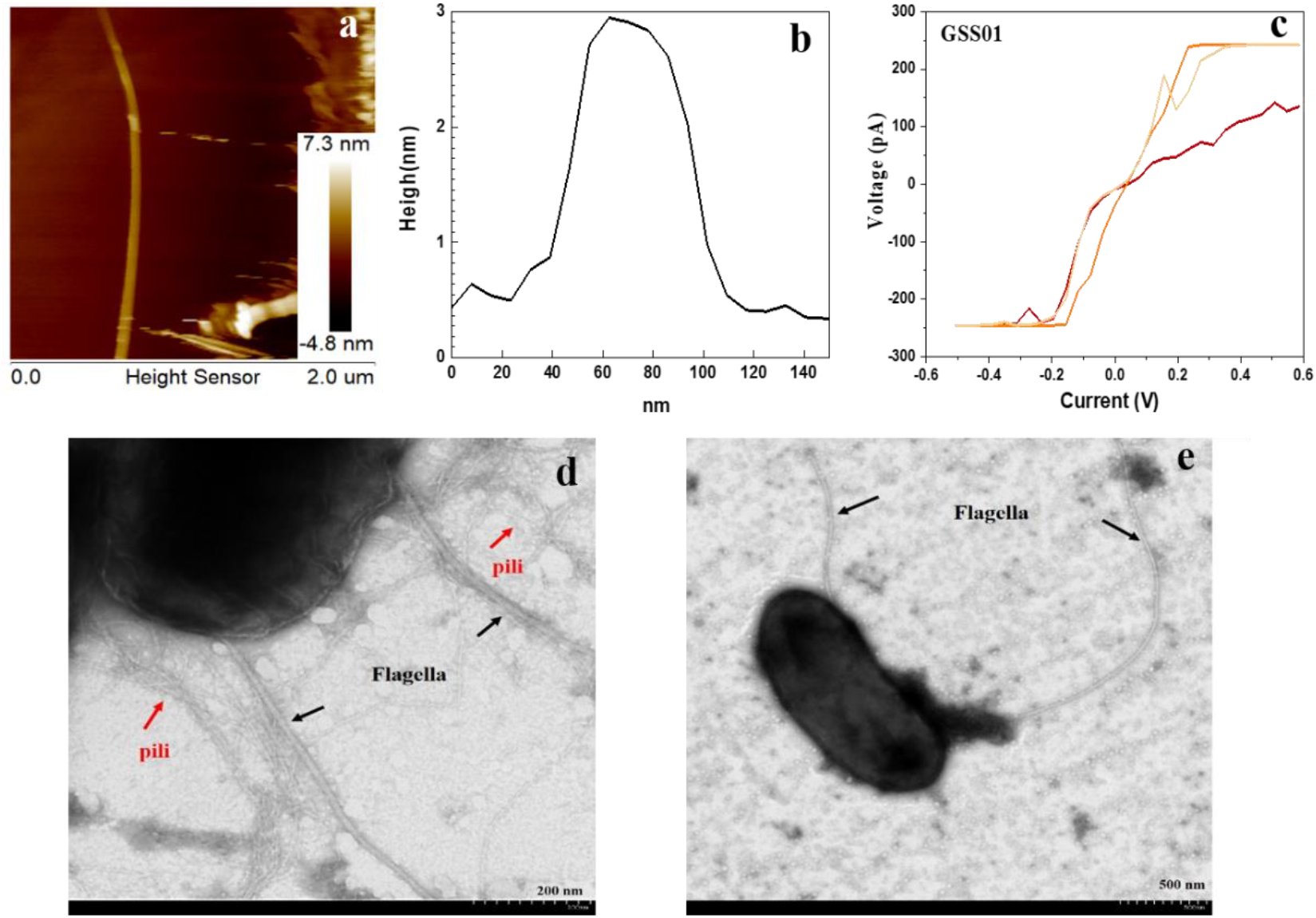
The characteristic of *G. soli* pili. (a) CP-AFM topography images of pili that purified from *G. soli* GSS01 (wild-type strain) deposited on HOPG surface. (b) The diameter of *G. soli* GSS01 pili. (c) Representative current-voltage (*I-V*) plots obtained after probing the transversal conductivity of *G. soli* GSS01 pili. (d) Transmission electron micrographs showing the presence of flagella and pili in *G. soli* GSS01. (e) Deletion of pilin gene of *G. soli* inhibited the grown of pili. Red arrows point toward the pili, while black arrows point toward the flagella.

To investigate the role of *G. soli* pili in extracellular electron transfer, the gene *pilA*, encoding the pilin protein, was deleted, which prevented the expression of lateral pili (Fig. 2d and 2e). The rate of soluble Fe(III) citrate reduction of the PilA-deficient mutant was similar to that of the wild-type strain (Fig. 3a), but the rate of insoluble Fe(III) oxide reduction decreased from 0.77 mM/day to 0.18 mM/day (Fig. 3b). Likewise, the PilA-deficient mutant also lost the capacity for electron transfer to an anode electrode (Fig. 3c) and formed a thinner biofilm than the wild-type strain (Fig. 3d and 3e).

**Fig. 3.**
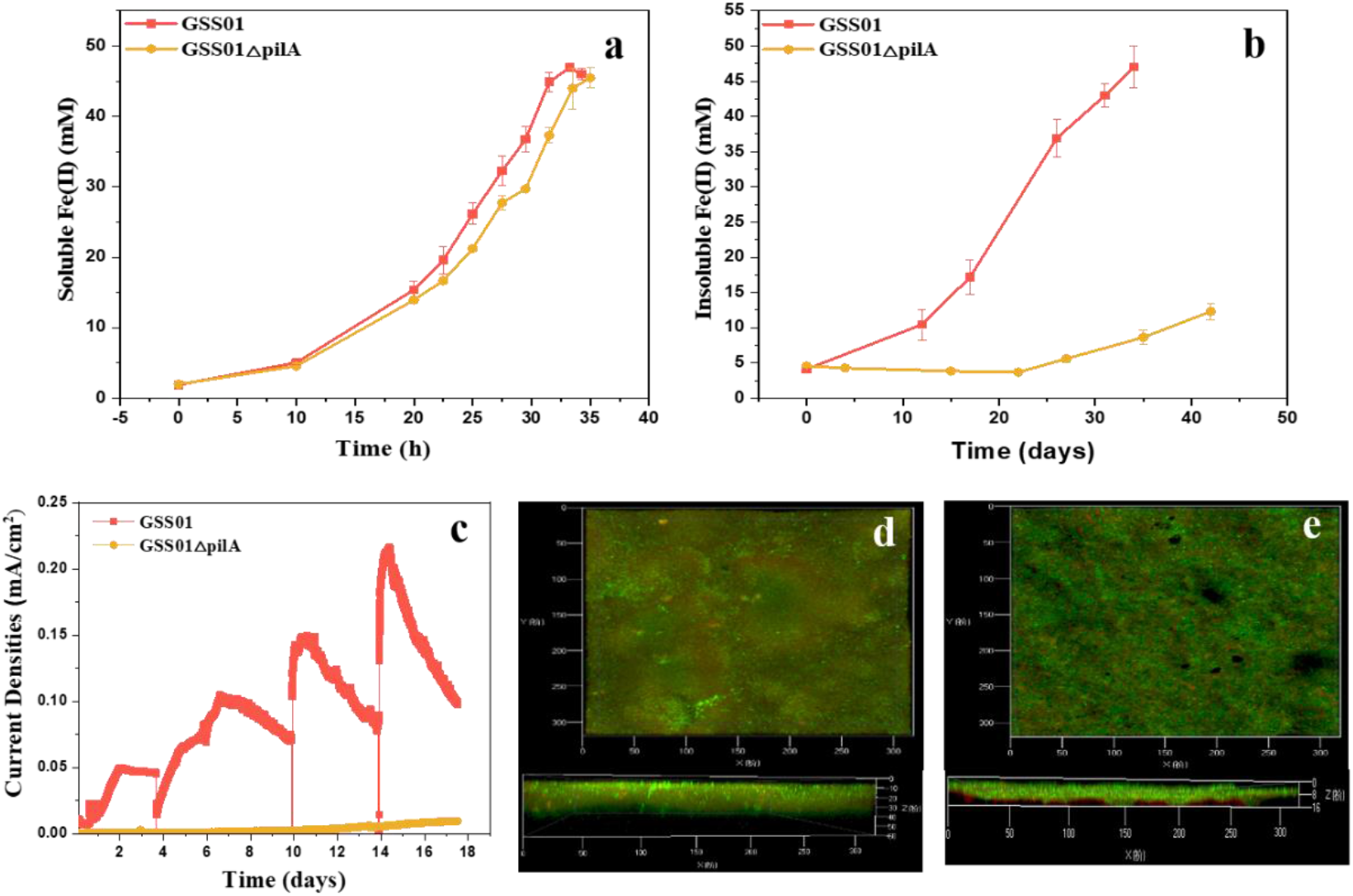
Fe(III) reduction and current production by *G. soli* pili-defective mutant. (a) The rate of Fe(III) citrate reduction of pili-defective mutant as well as that of wild-type strain. (b) The poorer rate of insoluble Fe(III) oxide reduction pili-defective mutant compared with wild-type strain. (c) Pili-defective mutant has lower current production than wild-type strain. The biofilm of pili-defective mutant (d) was thinner than wild-type strain (e).

### Heterologously expressing the pilin gene of *G. soli* in *G. sulfurreducens* yielded strain GSP, which produced abundant pili but was deficient in Fe(III) oxide reduction and current production

The *pilA* gene of *G. sulfurreducens* was replaced with the *pilA* gene of *G. soli* following previously described methods that have successfully approached the heterologous expression of *pilA* genes from other microorganisms in *G. sulfurreducens* (16, 17, 23, 24). Heterologously expressing the *pilA* gene of *G. soli* in *G. sulfurreducens* yielded a *G. sulfurreducens* strain, designated strain GSP (for *<u>G</u>. <u>s</u>oli* <u>p</u>ili). Strain GSP produced abundant pili (Fig. 4 and Fig. S2) with 4.4-fold lower conductivity than its host (Fig. 5). The difference in conductivity of pili between strain GSP and both wild-type *G. soli* and *G. sulfurreducens* was smaller and less than one order of magnitude, which suggests that their conductivity was comparable. The *pilA* transcription of strain GSP was analyzed, which further confirmed the expression of heterologous pili (supplementary Fig. S3). Strain GSP reduced Fe(III) citrate at a similar rate to the control strain, but the capability of reducing insoluble Fe(III) oxide was severely damaged. The data showed that the control strain produced Fe(II) at a rate of 0.27 mM/day, while the rate decreased to 0.04 mM/day for strain GSP (Fig. 6a and 6b). Similarly, strain GSP produced a current density that decreased by 10-fold and formed thinner biofilms than the control strain (Fig. 6c, d and e). The extracellular protein and extracellular cytochrome content of extracellular polymeric substances (EPS) extracted from the anode biofilm of strain GSP was significantly lower than that of the control strain (Fig. 7a and 7b), which corresponded to a lower current production and thinner biofilm (Fig. 6c and 6e).

**Fig. 4.**
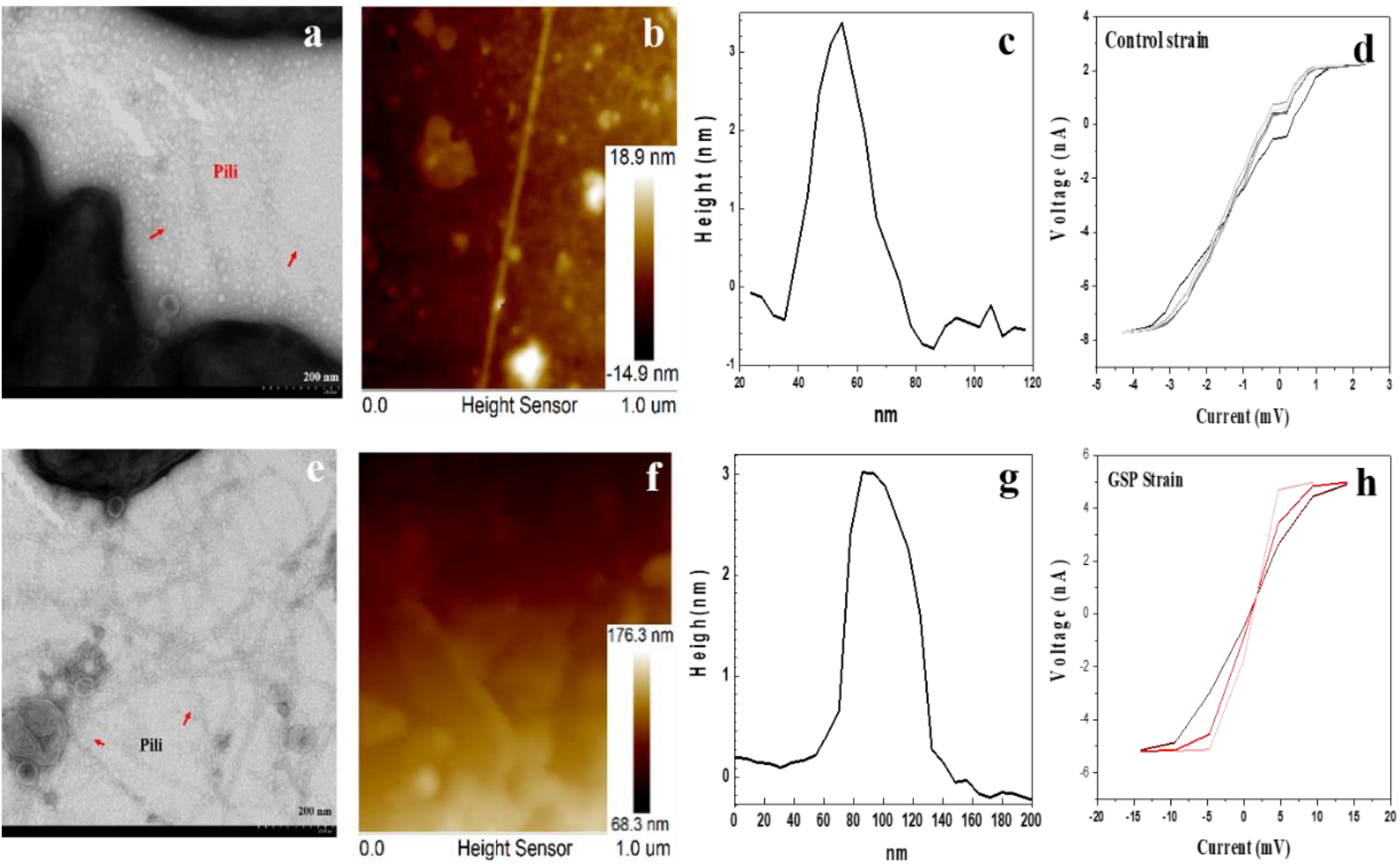
The characteristic of control strain and strain GSP pili. Transmission electron micrographs of negatively stained control strain (a) and strain GSP (e). CP-AFM topography images of pili purified from control strain (b) and strain GSP (f) deposited on a HOPG surface. The diameter of control strain (c) and strain GSP pili (g). Representative current-voltage (*I-V*) plots obtained after probing the transversal conductivity of control strain (d) and strain GSP pili (h).

**Fig. 5.**
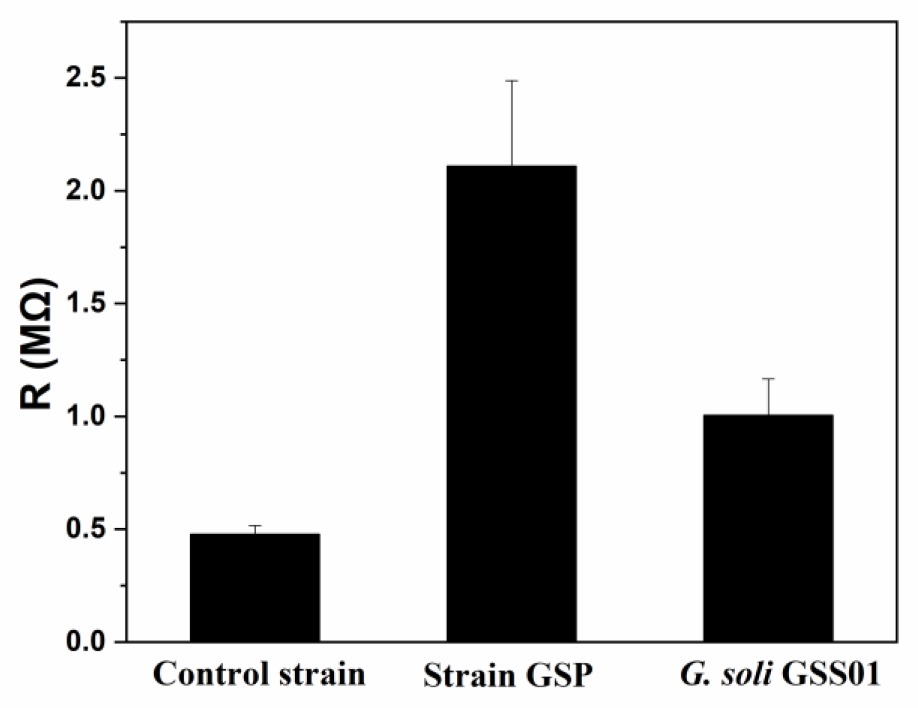
The resistances of pili obtained from *G. sulfurreducens* control strain, strain GSP and *G. soli* GSS01 (wild-type strain).

**Fig. 6.**
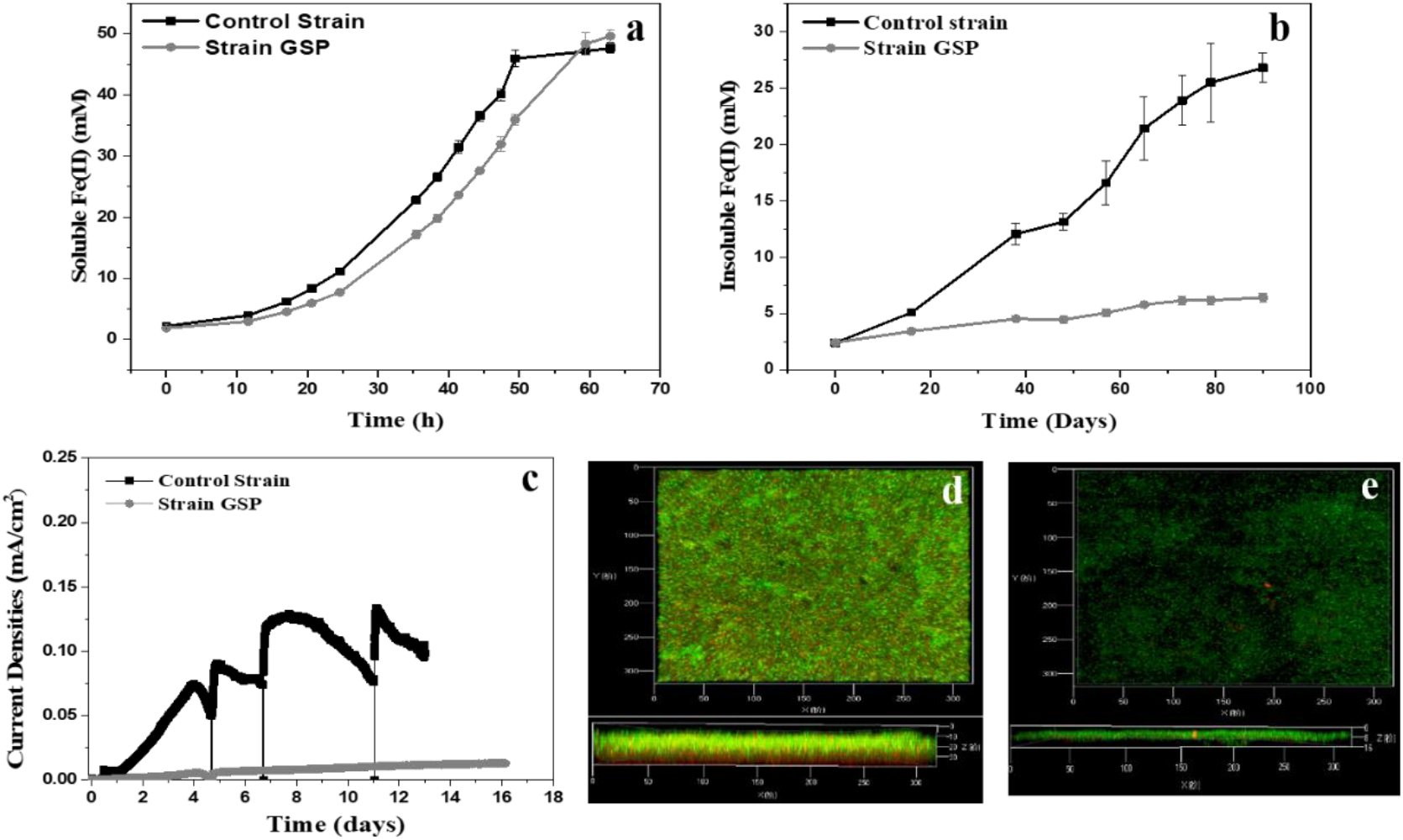
Fe(III) reduction and current production in control strain and strain GSP. (a) Both strain GSP and control strain have the similar rate of Fe(III) reduction when soluble Fe(III) citrate as the electron acceptor. (b) The rate of Fe(III) reduction of strain GSP was injured compared with control strain when insoluble Fe(III) oxide as the electron acceptor. (c) Current densities of strain GSP was lower than control strain. The biofilm of strain GSP (d) was thinner than the control strain (e).

**Fig. 7.**
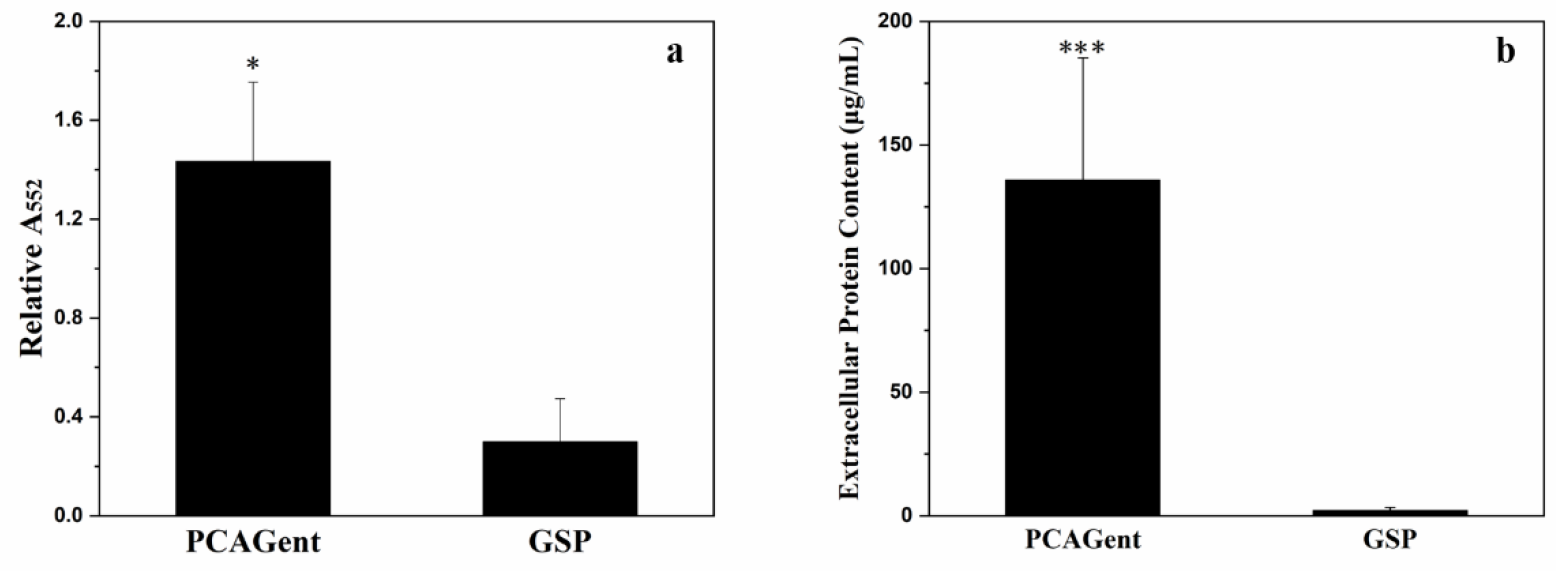
Relative cytochrome (a) and protein content (b) in EPS of biofilm extracts in the strain GSP and control strain (strain PCAGent). The results were shown as average ± SD (n=3) Significant differences in cytochrome and protein content relative to the control strain were identified in variance analysis (*P<0.05, **P<0.01, ***P< 0.001).

### Heterologous pili of *G. sulfurreducens* altered the content of outer-surface *c*-type cytochromes

Previous studies have demonstrated that heterologously expressing *pilA* in *G. sulfurreducens* did not alter the distribution of extracellular cytochromes (24), consistent with our results in the present study (Fig. 8). However, several bands from strain GSP exhibited slightly different stain intensities compared with the wild-type strain. Outer-surface *c*-type cytochrome OmcS showed distinct staining intensity. Two *trans*-outer membrane porin-cytochrome protein complexes of OmcB and OmcC, which are responsible for transferring electrons across the outer membrane to Fe(III) oxide, showed slight staining intensity (15, 25).

**Fig. 8.**
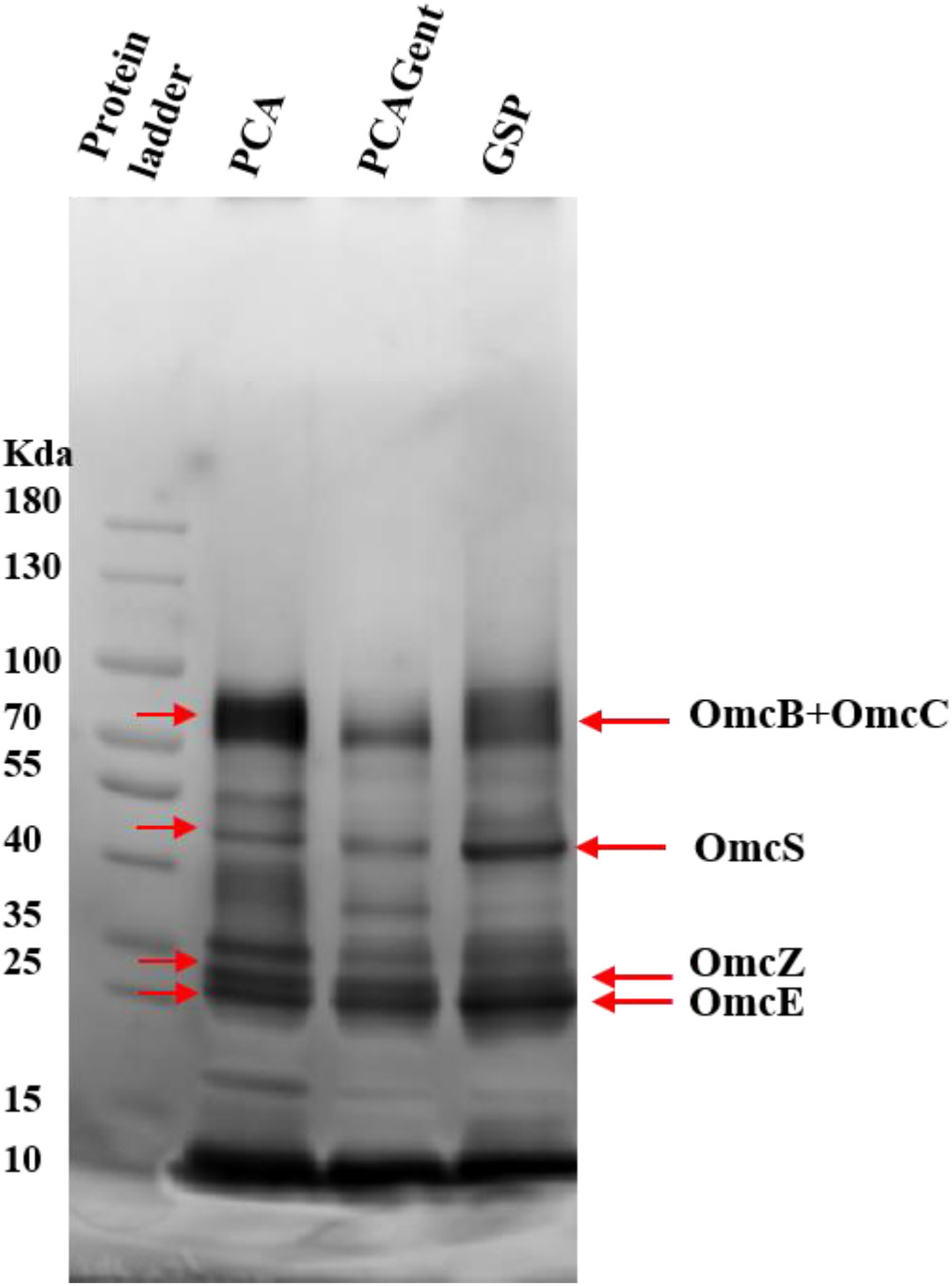
Extracellular cytochromes profile. Heme-stained SDS-PAGE of extracellular *c*-type cytochromes prepared from *G. sulfurreducens* PCA, control strain (strain PCAGent), and strain GSP. The localization of outer membrane *c*-type were labeled (red arrows) based on expected molecular weight (8 μg of protein was loaded into each lane).

The transcript abundance of the outer-surface *c*-type cytochrome gene of strain GSP changed significantly compared with the wild-type strain (Fig. 9). The abundance of *omcS* transcripts increased by two-fold, consistent with increased expression of OmcS (Fig. 8). OmcE, with 62.6% amino acid sequence identity to OmcS, remained the similar transcript abundance. Both OmcS and OmcE are known as key components in Fe(III) oxides and Mn(IV) oxides reduction (14). On the other hand, the abundance of *omcB* and *omcZ* transcripts was reduced 48.4 and 63.3 times, which involved in Fe(III) oxides reduction and current production (13, 15). The changes in the expression levels of *omcB* and *omcZ* corresponded to their transcription abundance (Fig. 8 and Fig. 9).

**Fig. 9.**
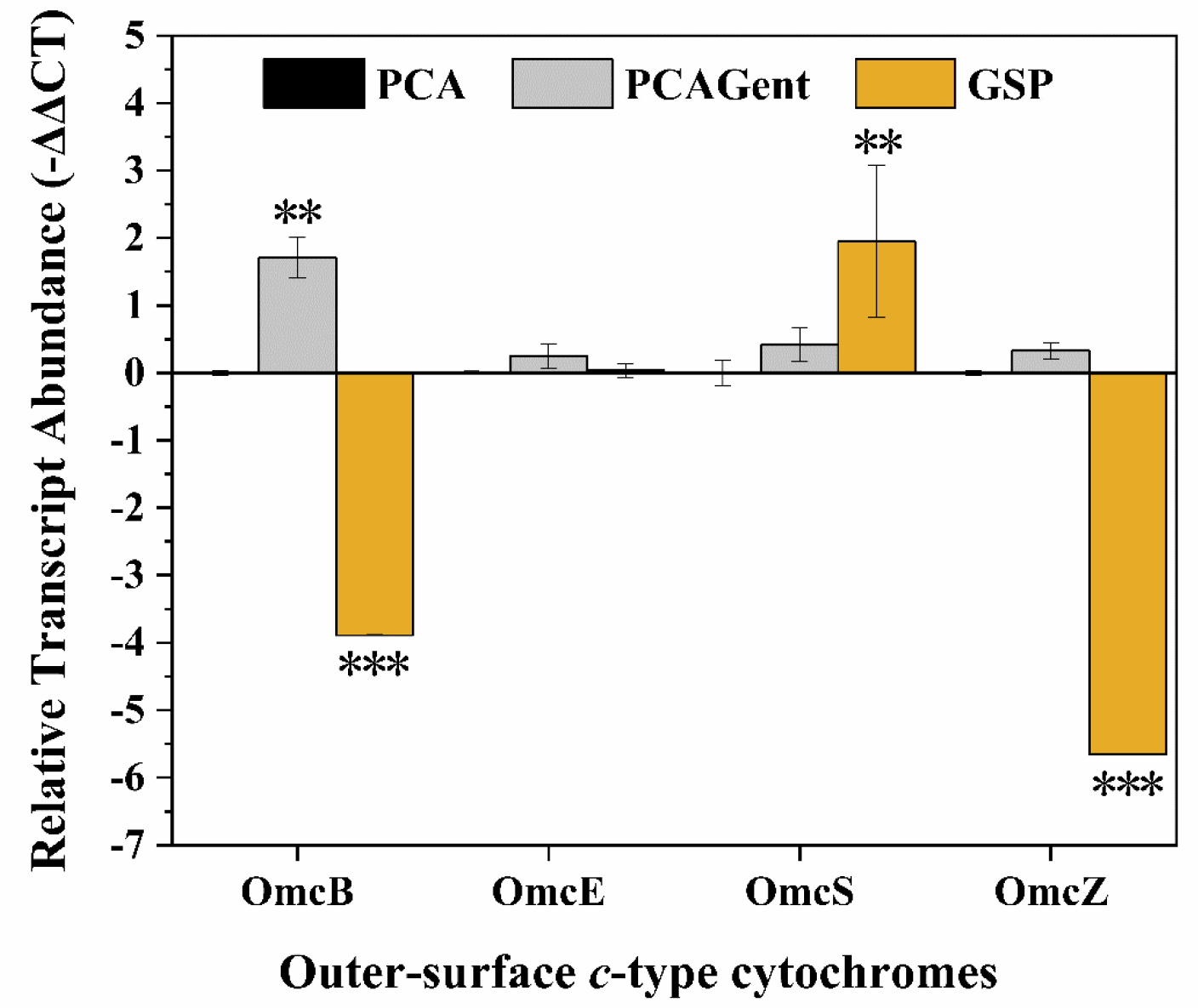
The transcript abundance of extracellular cytochromes (*OmcB, OmcE, OmcS* and *OmcZ*) from *G. sulfurreducens* control strain (strain PCAGent) (gray), strain GSP (yellow) compared to wild-type strain (black) in late-log NBAF medium. The housekeeping gene *proC* was used as internal control. Data are shown as average ± SD (n=4). Statistically significant changes relative to wild-type strain values were identified in variance analysis (*P < 0.05, **P < 0.01, ***P < 0.001).

## Discussion

In general, there are three procedures to verify the e-pili of diverse microorganisms: (i) determining the protein composition of the filaments (2); (ii) proving that the filaments are required for extracellular electron transfer (16); and (iii) documenting the conductivity of the filaments under physiologically relevant conditions (2). *G. soli*, closely related to *G. sulfurreducens*, has a similar pilin gene as *G. sulfurreducens* that encodes e-pili (22). The direct deletion of the pilin gene of *G. soli* demonstrated that *G. soli* pili is required for electron transport to Fe(III) oxides and electrodes, which plays an important role in extracellular electron transfer, consistent with *G. sulfurreducens* and *G. metallireducens* (4, 5, 9). Direct measurement showed that the conductivity of *G. soli* pili is similar to that of *G. sulfurreducens* pili (Fig. 1 and Fig. 5), which suggested that *G. soli* pili is sufficiently conductive along its length (10). The above results proved that *G. soli* has e-pili, corresponding to the definition, which is the next model of e-pili after the studies of *G. sulfurreducens* and *G. metallireducens*. It provides an additional model to promote studies of the role of pili in extracellular electron transfer and expands the range of electronic materials in biological devices.

An effective protocol for constructing gene deletions in *G. metallireducens* was modified for *G. soli* (9), including the high concentration of linear DNA for electroporation. The development of genetic manipulation of *G. soli* lays the foundation of technology for further studies of its physiological characteristics and is an important step in understanding the physiology of the genus *Geobacter*. For example, *G. soli* may act as an electron donating partner to connect other species, especially methanogens, by direct interspecies electron transfer because of their similar feature of extracellular electron transfer in both *G. sulfurreducens* and *G. metallireducens* (21).

*G. sulfurreducens* Strain GSP expressing the *G. soli* pilin gene, produced abundant pili (Fig. 4) with a conductivity comparable to the control strain of *G. sulfurreducens* expressing its native pilin gene, which was consistent with *G. soli* according to direct measurements (Fig. 5). These results demonstrate that the pilin gene of *G. soli* can properly assemble into pili when heterologously expressed in *G. sulfurreducens*, and the similar conductivity of pili between *G. soli* and *G. sulfurreducens* corresponds to their similar content of aromatic amino acids. It has been suggested that heterologously expressing pilin genes in *G. sulfurreducens* is a productive approach to yield pili, whose conductivity depends on the densities of aromatic amino acids of the pilin genes of the target microorganism (17). For example, introducing more aromatic rings of synthetic pilin monomers in *G. sulfurreducens* yields pili with higher conductivity than its host (26). In contrast, when the key aromatic amino acids of e-pili were substituted for alanine, the decreased content of aromatic amino acids of the pilin gene resulted in poor conductivity of pili (27). Additionally, the *G. sulfurreducens* strain heterologously expressing the pili monomer of *Desulfofervidus auxilii*, which has a lower density of aromatic amino acids than the native e-pili, produced pili with low conductivity (16).

Surprisingly, *G. sulfurreducens* strain GSP, heterologously expressing the pilin gene of *G. soli*, showed inhibited Fe(III) oxide reduction and electron transfer to an electrode. In addition to e-pili being essential for extracellular electron transfer (28, 29), the content (30) and composition of *c*-type cytochromes (13–15, 31) are also important for extracellular electron transfer. For example, the outer-surface *c*-type cytochrome OmcZ accumulates at the biofilm-electrode interface and plays a role as an electrochemical gate to facilitate the production of a high-density current and the formation of a thick biofilm (13, 32), and the outer-membrane *c*-type cytochrome OmcB is required for electron transfer to Fe(III) oxides (15). Previous studies have found that pili of *G. sulfurreducens*, including heterologous pili, are closely related to the expression of extracellular cytochromes (24, 29, 33). Direct evidence has shown that the deletion of *G. sulfurreducens* e-pili alters the distribution and composition of extracellular cytochromes (29, 33). Additionally, *G. sulfurreducens* e-pili is important for the localization of cytochromes, such as the outer-surface cytochrome OmcS (34), and the organization and anchoring of cytochromes to form thick biofilms and produce high currents (29, 35). Therefore, the results of further investigation on the expression of the outer-surface *c*-type cytochromes of strain GSP indicated that the heterologous pili of *G. sulfurreducens* reduced the content of outer-surface *c*-type cytochromes and consequently decreased Fe(III) oxide reduction and current production, which suggested that the content of outer-surface *c*-type cytochromes should be further researched when heterologous pili are expressed in *G. sulfurreducens*.

A number of outer membrane cytochromes are anchored to the extracellular polysaccharide network (36), particularly OmcZ, which is loosely bound to the extracellular matrix (32). Localization of type IV pili has been found to be correlated with extracellular polysaccharide in *Myxococcus xanthus*. The deletion of *pilA*, which encodes the type IV pilin of *M. xanthus*, prevented extracellular polysaccharide production (37, 38). Native PilA was not detected in a xapD-deficient strain of *G. sulfurreducens* with decreased expression of extracellular polysaccharide (36, 39). Thus, it can be speculated that PilA regulates the expression and colonization of extracellular polysaccharides, which affects the expression level of extracellular cytochromes. The low content of extracellular polymeric substances in strain GSP might explain the decrease in the expression level of outer membrane *c*-type cytochromes (Fig. 7a and 7b).

It was speculated that heterologous pili of *G. sulfurreducens* affected pilin-related proteins, which have an impact on the transcript abundance of outer-surface *c*-type cytochromes. *pilR*, the gene upstream of *pilA*, was knocked out, which resulted in downregulated transcript levels of *omcB* and *omcC* but upregulated transcript levels of *omcS* compared with wild-type strain (40). The type IV pili of *G. sulfurreducens* is considered type II secretions systems capable of translocating cytochromes from periplasm across the outer membrane (4, 41). The replacement of the native *G. sulfurreducens* pili with *G. soli* pili might influence the secretion and organization of outer-surface *c*-type cytochromes and consequently reduce the expression level of several extracellular cytochromes.

Heterologous expression of pilin genes from difficult-to-culture microorganisms offers a simple screening strategy to evaluate pili involved in extracellular electron transfer (16). The requirement of e-pili in *G. sulfurreducens* for the production of high current densities provides a clear phenotype to test the hypothesis that the heterologous pili gene encodes e-pili (16). However, the results of evaluating whether *G. soli* pili is electrically conductive by direct deletion and heterologous expression were different, which suggests that caution should be taken to determine whether e-pili of the target microorganism mediate extracellular electron transfer when a *G. sulfurreducens* strain with heterologously expressed pili produces low current densities. For example, the pilin gene of *Desulfofervidus auxilii* was heterologously expressed in *G. sulfurreducens* to yield a strain that produced low current densities with relatively thin biofilms, and the analysis suggested that the hypothesis that the sulfate reducer wires itself to ANME-1 microbes with e-pili be reevaluated (16). However, it is not rigorous to assume that the low current densities of this strain resulted from the lack of e-pili because *G. sulfurreducens* has diverse electron transfer routes for current production (42), and further research on *c*-type cytochromes (32) is lacking. After all, pili of the sulfate reducer was highly expressed in syntrophic consortia (43). Therefore, it is insufficient to evaluate whether *D. auxilii* pili mediate direct electron transfer to its partner ANME-1 (16). This suggested that the method of heterologously expressing pilin genes of phylogenetically diverse microorganisms in *G. sulfurreducens* to screen e-pili via evaluation of high current densities should be revisited.

This study found that *G. soli* pili is electrically conductive and participate in Fe(III) oxide reduction and electron transfer to an electrode, as previously reported in *G. sulfurreducens* and *G. metallireducens* (9, 44). The development of genetic manipulation of *G. soli* is an important strategy to further understand the physiology of *Geobacter* species, which play an important role in anaerobic environments. *G. sulfurreducens* strain GSP, which heterologously expressing the e-pili of *G. soli* in *G. sulfurreducens*, is deficient in Fe(III) oxide reduction and current production due to the impaired content of outer surface *c*-type cytochromes. The analysis demonstrated that the heterologous pili of *G. sulfurreducens* reduced the content of *c*-type cytochromes, which suggested that the content of extracellular cytochromes should be considered, when employing *G. sulfurreducens* to heterologously express pilin genes, especially for screening e-pili from diverse microorganisms.

## Materials and methods

### Bacterial strains, plasmids and cultivation conditions

All bacterial strains and plasmids used in this study are summarized in Table S1 in the supplemental material. The wild-type strain *G. sulfurreducens* PCA (DSM 12127) was purchased from the German Collection of Microorganisms and Cell Culture. *G. soli* GSS01 was provided by Zhou’s lab (20). Both strains were cultured at 30 °C under strict anaerobic conditions (80%/20%; N2/CO2) and used for the construction of related mutants. *G. sulfurreducens* was routinely cultured in mineral-based medium containing acetate (15 mM) as the electron donor and fumarate (40 mM) as the electron acceptor, as previously described (45). *G. soli* was cultured in freshwater medium (46) containing Fe(III) citrate (56 mM) and acetate (16.5 mM) as the electron acceptor and donor, respectively. Chemically competent *Escherichia coli* DH5α cells (Takara, Daliang, China) were regularly used for cloning and were cultured at 37 °C in Luria-Bertani medium supplemented with the appropriate antibiotics.

### Deletion of PilA from *G. soli*

Primers used to construct the mutants and validate the genotypes are listed in Table S2. Deletion by single-step gene replacement of *pilA* is summarized in Figure S6. Construction of the *pilA::Emr^r^* allele was performed following a protocol previously described (13). The erythromycin resistance cassette flanked by *loxP* sites was constructed as follows: the primer pair gmf/gmr was used to amplify the gentamycin resistance cassette flanked by *loxp* sites from the plasmid pCM351 (33). The PCR products were TA cloned into pMD™19, and then the obtained plasmid, pMD19-GmloxP, was digested with *Mlu* I to excise the gentamycin resistance gene with its promoter. The erythromycin resistance gene was amplified from plasmid pMG36e using primer pair EmrF/EmrR and then connected with linearized pMD19-GmloxP using In-Fusion HD Enzyme (Takara, Daliang, China), generating the plasmid pMD19-EmrloxP. The primer pair gmf/gmr was also used to amplify *loxP*-flanked erythromycin resistance genes from the plasmid pMD19-EmrloxP. The primer pair GS01PilAUPF/GS01PilAUPR was used to amplify ≈500 bp upstream of *pilA*, and GS01PilADNF/GS01PilADNR was used to amplify ≈500 bp downstream of *pilA* with GSS01 genomic DNA as a template. The two fragments plus the *loxP*-flanked erythromycin resistance gene were connected with the linear plasmid pUC19 using In-Fusion HD Enzyme (Takara, Daliang, China), generating the plasmid pUC-GSS01PilA. The sequence of the construction was verified by Sanger sequencing and then linearized with ScaI (NEB, MA, USA).

*G. soli* was made electrocompetent following a protocol previously described (9). Icecold freshly prepared competent cells (25 μl) were mixed with 4.5 μg of linear DNA and transferred to 0.1 cm-gap electroporation cuvettes (Bio-Rad, CA, USA). A pulse of 1.5 kV was applied to the cuvette with a MicroPulser electroporation apparatus (Bio-Rad, CA, USA). Immediately following electroporation, cells were transferred to an anaerobic pressure tube containing 10 ml of acetate-Fe(III) citrate medium with ferrous ammonium sulfate and yeast extract. The electroporated cells were allowed to recover for 18 h at 30 °C and then spread on citrate Fe(III) solid plates supplemented with 200 μg/ml erythromycin.

A single colony was selected then verified using the primer pair VerEmrF/VerGS01PilAR to confirm that the erythromycin-resistant gene was integrated into the GSS01 chromosome (Fig. S2). The primer pair GS01PilAF-QPCR/GS01PilAR-QPCR was used to confirm that the coding sequence of *pilA* was deleted from the GSS01 chromosome. The primer pair VerpUC19F/VerpUC19R was used to confirm that there was no circular plasmid in the GSS01 chromosome.

### Heterologously expressing e-pili of *G. soli* in *G. sulfurreducens*

Strain GSP was constructed from *G. sulfurreducens* by using a previously described approach for the expression of heterologous *pilA* genes (Fig. S6) (24). The primer pair GSU1495UPF/GSU1495UPR was used to amplify 510 bp upstream of GSU1496 with the PCA chromosome serving as the template to generate fragment 1. The primer pair PilASNPDNF/PilASNPDNR amplified the promoter region of the *G*. *sulfurreducens pilA* gene to generate fragment 2. For the generation of fragment 3, the primer pair GSPilACF/GSPilACR was used to amplify 555 bp downstream of GSU1496. The primer pair GS01PilAF/GS01PilAR amplified *pilA* (locus tag SE37_07695) with GSS01 genomic DNA as the template to generate fragment 4. The above four fragments plus the fragment of the gentamycin resistance cassette flanked by *loxp* sites were connected and inserted into the linearized plasmid pUC19 using the In-Fusion Cloning Kit (TaKaRa, Daliang, China), generating the plasmid pUC-GSP. The plasmid was verified by Sanger sequencing, linearized with ScaI (NEB, MA, USA), and electroporated into electrocompetent *G. sulfurreducens* as previously described (45).

The primer pair gmm/GS01PilAR was used to verify the gentamycin resistance gene in the positive recombinant strains (Fig. S2). The primer pair PilAf/PilAr was used to confirm that the *pilA* gene of *G. soli* was replaced with the *pilA* gene of *G. sulfurreducens*. Finally, the primer pair gmm. F1/GS01PilAR was used to further verify the recombination of the *pilA* gene of *G. soli* in the mutant strains. The control strain containing the gentamycin resistance cassette flanked by *loxp* sites was constructed as follows. The primer pair PilASNPDNF/GSU1497SNPDNR was used to amplify GSU1496 and upstream of GSU1497. The PCR product was combined with fragment 2 and the gentamycin resistance cassette flanked by *loxp* sites and then inserted into the linearized plasmid pUC19 by an In-Fusion Cloning Kit (TaKaRa, Daliang, China) to generate the plasmid pUC19-Gent. The plasmid was linearized using ScaI and transferred to competent cells for electroporation as previously described (45).

The primer pair gmm/PilASNPDNR was used to verify the gentamycin resistance gene in mutant strains, while the wild-type strain had no product. The primer pair PilAf/PilAr was used as a positive control in the control strain and the wild-type strain. All mutant strains were verified by PCR (Fig. S2) and Sanger sequencing.

### RNA extraction and RT-qPCR

All strains were grown to the late-log stage in NBAF medium with the same OD_600_ as previously described (45). Cells were harvested by centrifugation and then frozen with liquid nitrogen. RNA was extracted with a Bacterial RNA Extraction Kit and was immediately reverse-transcribed into cDNA using a PrimeScriptT™ RT-PCR Kit (TAKARA, Daliang, China). The common primer pair GSPilACF/GSPilACR was used to test genomic DNA contamination. The primer pair PilAf/PilAr was used to identify the transcription of pilin in the wild-type and control strains, and the primer pair GS01PilAF-QPCR/GS01PilAR-QPCR was used to test the transcription of recombinant pilin in the GSP strain. PCR products and extracted RNA were analyzed on a 1.5% agarose/TAE gel (Fig. S3).

Transcripts for key extracellular *c*-type cytochromes were quantified by reverse transcription-quantitative PCR (RT-qPCR). The gene targets were outer membrane *c*-type cytochromes *omcB* (GSU2737), *omcE* (GSU0618), *omcS* (GSU2504) and *omcZ* (GSU2078). The housekeeping gene *proC* was used as a control as previously described (13). RT-qPCR was performed using CFX Connect (Bio-Rad, CA, USA) and the primers listed in Table S2. The relative fold change of expression for each target gene versus the *proC* internal control for each strain versus the wild-type strain was then calculated with the formula 2^−ΔΔCT^ for four replicate samples. Statistically significant changes in gene expression relative to the control gene were determined by analysis of variance.

### TEM and CSLM

Anode-grown cells were examined with transmission electron microscopy. Samples were negatively stained with 2% uranyl acetate and imaged by a HITACHI H-7650 transmission electron microscope at an accelerating voltage of 80 kV. The anode biofilms were rinsed with freshwater medium and stained with a LIVE/DEAD BacLight Bacterial Viability Kit (Thermo Fisher Scientific, NY, USA) when the currents reached the maximum value (47). A confocal scanning laser microscope (ZEISS LSM T-PMT) with a 20× objective lens was used for imaging.

### Pili preparation and conductivity measurements by CP-AFM

Pili from the anode biofilm of wild-type *G. soli*, wild-type *G. sulfurreducens*, and strain GSP were purified in ethanolamine buffer by two centrifugations as previously described (17). The final pili preparation was resuspended in ethanolamine buffer and stored at 4 °C. The pili in ethanolamine buffer were deposited on the surface of freshly cleaved highly oriented pyrolytic graphite (HOPG) to probe the transversal conductivity of hydrated pili by conductive probe-atomic force microscopy (CP-AFM) as previously described (29, 48). HOPG coated with the samples was electrically connected to a CP-AFM system (Fastscan, Bruker, German) with silver paint. The CP-AFM tip was used for scanning and imaging the pili of individual samples in the peak force tapping mode (Fig. S4). The transversal conductivity of different points along each pilus filament was measured by the CP-AFM tip. At least three current-voltage (*I-V*) curves were obtained at different points while applying a bias voltage within the ± 1 V range (10 nN force, 0.5 Hz rate). The electrical resistance (*R*) at different points of each pilus was calculated from the linear portion of each *I-V* curve, as described elsewhere (49), and the average and s.d. of all the resistance values were used to compare the conductivity of the pili from the different strains.

### Current production, Fe(III) reduction and Heme-staining of SDS-PAGE

A two-chambered H-cell three-electrode system with acetate (15 mM) as the electron donor and graphite plate (15 cm^2^) as the electron acceptor was used to characterize the current production of each strain (13). The system was connected with a potentiostat (CH Instruments Ins., Shanghai, China) to record the current data, and the working electrode was poised at +300 mV versus Ag/AgCl. The different strains were grown to mid-log and inoculated into the working chamber; then, the systems were operated for a total of four cycles in sequencing batch mode (9).

For Fe(III) reduction studies, Fe(III) citrate (56 mM) or synthetic ferrihydrite (100 mM) served as the sole electron acceptor, and acetate (15 mM) served as the electron donor. Fe(II) production was measured with a ferrozine assay (50).

Extracellular proteins were collected from late-log cells grown in NBAF medium as previously described (14) and quantified using the Pierce™ BCA Protein Assay Kit (Thermo Fisher Scientific, MA, USA). The amount of non-reducing loading buffer was used to equal the amount of protein. The samples were loaded on 4-20% BeyGel™ Plus PAGE (Beyotime, Shanghai, China) and heme-stained (51).

### Assays for extracellular cytochrome content in extracted EPS of biofilm

The EPS of biofilms were extracted using a previously modified method (52, 53). The biofilm was gently scraped off the graphite plate and dissolved in 3 ml of 0.9% (w/v) NaCl solution. The sample was vortexed in a tube for 5 min at setting 10 to disperse the cells. Then, an equal volume of 2% Na2-EDTA (53.7 mM, pH 7.0, in 0.9% NaCl) was mixed with cells at 4 °C for 3 h and centrifuged at 5,000 g for 20 min. The supernatant was filtered through a 0.22 μm membrane filter.

The EPS of *Geobacter* biofilms contain abundant extracellular cytochromes, and the reduced extracellular cytochromes represent the ability to deliver electrons to extracellular electron acceptors. The extracellular cytochrome content of EPS was determined using ultraviolet-visible spectroscopy (UV-2600, Shimadzu, Japan) from 350 nm to 650 nm as previously described (29). Reduced spectra at 552 nm after adding dithionite as a reducing agent anaerobically were used to estimate the extracellular cytochrome content in extracted EPS of biofilms (54) (Fig. S5). Statistically significant differences were determined using the variance test for triplicate cultures.

## Acknowledgments

The research was supported by the National Program for Support of Top-Notch Young Professionals, Guangdong Special Support Program (2017TX04Z351) and Science and Technology Program of Guangzhou, China (201903010071).

## References

1. Lovley DR, Ueki T, Zhang T, Malvankar NS, Shrestha PM, Flanagan KA, Aklujkar M, Butler JE, Giloteaux L, Rotaru A-E, Holmes DE, Franks AE, Orellana R, Risso C, Nevin KP. 2011. *Geobacter:* the microbe electric’s physiology, ecology, and practical applications. Adv Microb Physiol 59:1–100.

2. Lovley DR. 2017. Electrically conductive pili: biological function and potential applications in electronics. Curr Opin Electrochem 4:190–198.

3. Shi L, Dong HL, Reguera G, Beyenal H, Lu AH, Liu J, Yu HQ, Fredrickson JK. 2016. Extracellular electron transfer mechanisms between microorganisms and minerals. Nat Rev Microbiol 14: 651–662.

4. Reguera G, McCarthy KD, Mehta T, Nicoll JS, Tuominen MT, Lovley DR. 2005. Extracellular electron transfer via microbial nanowires. Nature 435:1098–1101.

5. Reguera G, Nevin KP, Nicoll JS, Covalla SF, Woodard TL, Lovley DR. 2006. Biofilm and nanowire production leads to increased current in *Geobacter sulfurreducens* fuel cells. Appl Environ Microbiol 72:7345–7348.

6. Rotaru AE, Shrestha PM, Liu F, Markovaite B, Chen S, Nevin KP, Lovley DR. 2014. Direct interspecies electron transfer between *Geobacter metallireducens* and *Methanosarcina barkeri*. Appl Environ Microbiol 80:4599–4605.

7. Rotaru A-E, Shrestha PM, Liu F, Shrestha M, Shrestha D, Embree M, Zengler K, Wardman C, Nevin KP, Lovley DR. 2014. A new model for electron flow during anaerobic digestion: direct interspecies electron transfer to *Methanosaeta* for the reduction of carbon dioxide to methane. Energy Environ Sci. 7:408–415.

8. Lovley DR. 2017. e-Biologics: Fabrication of Sustainable Electronics with “Green” Biological Materials. mBio 8:e00695–17.

9. Tremblay PL, Aklujkar M, Leang C, Nevin KP, Lovley D. 2012. A genetic system for *Geobacter metallireducens*: role of the flagellin and pilin in the reduction of Fe(III) oxide. Environ Microbiol Rep 4:82–88.

10. Adhikari RY, Malvankar NS, Tuominen MT, Lovley DR. 2016. Conductivity of individual *Geobacter* pili. RSC Adv 6:8354–8357.

11. Malvankar NS, Vargas M, Nevin K, Tremblay PL, Evans-Lutterodt K, Nykypanchuk D, Martz E, Tuominen MT, Lovley DR. 2015. Structural basis for metallic-like conductivity in microbial nanowires. mBio 6:e00084–15.

12. Smith JA, Lovley DR, Tremblay P-L. 2013. Outer Cell Surface Components Essential for Fe(III) Oxide Reduction by *Geobacter metallireducens*. Appl Environ Microbiol 79:901–907.

13. Nevin KP, Kim BC, Glaven RH, Johnson JP, Woodard TL, Methe BA, Didonato RJ, Covalla SF, Franks AE, Liu A, Lovley DR. 2009. Anode biofilm transcriptomics reveals outer surface components essential for high density current production in *Geobacter sulfurreducens* fuel cells. PLoS One 4:e5628.

14. Mehta T, Coppi MV, Childers SE, Lovley DR. 2005. Outer membrane c-type cytochromes required for Fe(III) and Mn(IV) oxide reduction in *Geobacter sulfurreducens*. Appl Environ Microbiol 71:8634–8641.

15. Leang C, Coppi MV, Lovley DR. 2003. OmcB, a c-type polyheme cytochrome, involved in Fe(III) reduction in *Geobacter sulfurreducens*. J Bacteriol 185:2096–2103.

16. Walker DJF, Adhikari RY, Holmes DE, Ward JE, Woodard TL, Nevin KP, Lovley DR. 2018. Electrically conductive pili from pilin genes of phylogenetically diverse microorganisms. ISME J 12:48–58.

17. Tan Y, Adhikari RY, Malvankar NS, Ward JE, Woodard TL, Nevin KP, Lovley DR. 2017. Expressing the *Geobacter metallireducens* PilA in *Geobacter sulfurreducens* yields pili with exceptional conductivity. mBio 8:e02203–16.

18. Caccavo F, Lonergan DJ, Lovley DR, Davis M, Stolz JF, Mclnerney MJ. 1994. *Geobacter sulfurreducens* sp. nov., a hydrogen-and acetate-oxidizing dissimilatory metal-reducing microorganism. Appl Environ Microbiol 60: 3752–3759.

19. Sun D, Wang A, Cheng S, Yates M, Logan BE. 2014. *Geobacter anodireducens* sp nov., an exoelectrogenic microbe in bioelectrochemical systems. Int J Syst Evol Microbiol 64:3485–3491.

20. Zhou S, Yang G, Lu Q, Wu M. 2014. *Geobacter soli* sp. nov., a dissimilatory Fe(III)-reducing bacterium isolated from forest soil. Int J Syst Evol Microbiol 64:3786–3791.

21. Cai X, Huang L, Yang G, Yu Z, Wen J, Zhou S. 2018. Transcriptomic, proteomic, and bioelectrochemical characterization of an exoelectrogen *Geobacter soli* grown with different electron acceptors. Front Microbiol 9:3111.

22. Holmes DE, Dang Y, Walker DJF, Lovley DR. 2016. The electrically conductive pili of *Geobacter* species are a recently evolved feature for extracellular electron transfer. Microb Genom 2:e000072.

23. Tan Y, Adhikari RY, Malvankar NS, Ward JE, Nevin KP, Woodard TL, Smith JA, Snoeyenbos-West OL, Franks AE, Tuominen MT, Lovley DR. 2016. The low conductivity of *Geobacter uraniireducens* pili suggests a diversity of extracellular electron transfer mechanisms in the genus *Geobacter*. Front Microbiol 7:980.

24. Liu X, Tremblay P-L, Malvankar NS, Nevin KP, Lovley DR, Vargas M. 2014. A *Geobacter sulfurreducens* strain expressing *Pseudomonas aeruginosa* type IV pili localizes OmcS on pili but is deficient in Fe(III) oxide reduction and current production. Appl Environ Microbiol 80:1219–1224.

25. Santos TC, Silva MA, Morgado L, Dantas JM, Salgueiro CA. 2015. Diving into the redox properties of *Geobacter sulfurreducens* cytochromes: a model for extracellular electron transfer. Dalton Trans 44:9335–9344.

26. Tan Y, Adhikari RY, Malvankar NS, Pi S, Ward JE, Woodard TL, Nevin KP, Xia Q, Tuominen MT, Lovley DR. 2016. Synthetic biological protein nanowires with high conductivity. Small 12:4481–4485.

27. Vargas M, Malvankar NS, Tremblay PL, Leang C, Smith JA, Patel P, Snoeyenbos-West O, Nevin KP, Lovley DR. 2013. Aromatic amino acids required for pili conductivity and long-range extracellular electron transport in *Geobacter sulfurreducens*. mBio 4:e00105–00113.

28. Ordonez MV, Schrott GD, Massazza DA, Busalmen JP. 2016. The relay network of *Geobacter* biofilms. Energy Environ Sci 9:2677–2681.

29. Steidl RJ, Lampa-Pastirk S, Reguera G. 2016. Mechanistic stratification in electroactive biofilms of *Geobacter sulfurreducens* mediated by pilus nanowires. Nat Commun 7:12217.

30. Estevez-Canales M, Kuzume A, Borjas Z, Fueg M, Lovley D, Wandlowski T, Esteve-Nunez A. 2015. A severe reduction in the cytochrome C content of *Geobacter sulfurreducens* eliminates its capacity for extracellular electron transfer. Environ Microbiol Rep 7:219–226.

31. Peng L, Zhang Y. 2017. Cytochrome OmcZ is essential for the current generation by *Geobacter sulfurreducens* under low electrode potential. Electrochim Acta 228:447–452.

32. Inoue K, Leang C, Franks AE, Woodard TL, Nevin KP, Lovley DR. 2011. Specific localization of the c-type cytochrome OmcZ at the anode surface in current-producing biofilms of *Geobacter sulfurreducens*. Environ Microbiol Rep 3:211-217.

33. Liu X, Zhuo S, Rensing C, Zhou S. 2018. Syntrophic growth with direct interspecies electron transfer between pili-free *Geobacter* species. ISME J 12:2142–2151.

34. Leang C, Qian X, Mester T, Lovley DR. 2010. Alignment of the c-type cytochrome OmcS along pili of *Geobacter sulfurreducens*. Appl Environ Microbiol 76:4080–4084.

35. Snider RM, Strycharz-Glaven SM, Tsoi SD, Erickson JS, Tender LM. 2012. Long-range electron transport in *Geobacter sulfurreducens* biofilms is redox gradient-driven. Proc Natl Acad Sci U S A 109:15467–15472.

36. Rollefson JB, Stephen CS, Tien M, Bond DR. 2011. Identification of an extracellular polysaccharide network essential for cytochrome anchoring and biofilm formation in *Geobacter sulfurreducens*. J Bacteriol 193:1023–1033.

37. Black WP, Xu Q, Yang Z. 2006. Type IV pili function upstream of the Dif chemotaxis pathway in *Myxococcus xanthus* EPS regulation. Mol Microbiol 61:447–456.

38. Yang Z, Lux R, Hu W, Hu C, Shi W. 2010. PilA localization affects extracellular polysaccharide production and fruiting body formation in *Myxococcus xanthus*. Mol Microbiol 76:1500–1513.

39. Flanagan KA, Leang C, Ward JE, Lovley DR. 2017. Improper localization of the OmcS cytochrome may explain the inability of the xapD-Deficient mutant of *Geobacter sulfurreducens* to reduce Fe(III) oxide. bioRxiv https://doi.org/10.1101/114900.

40. Juarez K, Kim BC, Nevin K, Olvera L, Reguera G, Lovley DR, Methe BA. 2009. PilR, a transcriptional regulator for pilin and other genes required for Fe(III) reduction in *Geobacter sulfurreducens*. J Mol Microbiol Biotechnol 16:146–158.

41. Richter LV, Sandler SJ, Weis RM. 2012. Two isoforms of *Geobacter sulfurreducens* PilA have distinct roles in pilus biogenesis, cytochrome localization, extracellular electron transfer, and biofilm formation. J Bacteriol 194:2551–2563.

42. Levar CE, Chan CH, Mehta-Kolte MG, Bond DR. 2014. An inner membrane cytochrome required only for reduction of high redox potential extracellular electron acceptors. mBio 5:e02034–14.

43. Wegener G, Krukenberg V, Riedel D, Tegetmeyer HE, Boetius A. 2015. Intercellular wiring enables electron transfer between methanotrophic archaea and bacteria. Nature 526:587–590.

44. Methe BA, Nelson KE, Eisen JA, Paulsen IT, Nelson W, Heidelberg JF, Wu D, Wu M, Ward N, Beanan MJ, Dodson RJ, Madupu R, Brinkac LM, Daugherty SC, DeBoy RT, Durkin AS, Gwinn M, Kolonay JF, Sullivan SA, Haft DH, Selengut J, Davidsen TM, Zafar N, White O, Tran B, Romero C, Forberger HA, Weidman J, Khouri H, Feldblyum TV, Utterback TR, Van Aken SE, Lovley DR, Fraser CM. 2003. Genome of *Geobacter sulfurreducens:* metal reduction in subsurface environments. Science 302:1967–1969.

45. Coppi MV, Leang C, Sandler SJ, Lovley DR. 2001. Development of a genetic system for *Geobacter sulfurreducens*. Appl Environ Microbiol 67:3180–3187.

46. Bond DR, Lovley DR. 2003. Electricity production by *Geobacter sulfurreducens* attached to electrodes. Appl Environ Microbiol 69:1548–1555.

47. Reardon PN, Mueller KT. 2013. Structure of the type IVa major pilin from the electrically conductive bacterial nanowires of *Geobacter sulfurreducens*. J Biol Chem 288:29260–29266.

48. Hu J, Zeng C, Liu G, Lu Y, Zhang R, Luo H. 2019. Enhanced sulfate reduction accompanied with electrically-conductive pili production in graphene oxide modified biocathodes. Bioresour Technol 282:425–432.

49. Lampa-Pastirk S, Veazey JP, Walsh KA, Feliciano GT, Steidl RJ, Tessmer SH, Reguera G. 2016. Thermally activated charge transport in microbial protein nanowires. Sci Rep 6:23517.

50. Anderson RT, Lovley DR. 2010. Naphthalene and benzene degradation under Fe(III)-reducing conditions in petroleum-contaminated aquifers. Bioremediat J 3:121–135.

51. Thomas PE, Ryan D, Levin W. 1976. An improved staining procedure for the detection of the peroxidase activity of cytochrome P-450 on sodium dodecyl sulfate polyacrylamide gels. Anal Biochem 75:168–176.

52. Dai YF, Xiao Y, Zhang EH, Liu LD, Qiu L, You LX, Dummi Mahadevan G, Chen BL, Zhao F. 2016. Effective methods for extracting extracellular polymeric substances from *Shewanella oneidensis* MR-1. Water Sci Technol 74:2987–2996.

53. Yang G, Lin J, Zeng EY, Zhuang L. 2019. Extraction and characterization of stratified extracellular polymeric substances in *Geobacter* biofilms. Bioresour Technol 276:119–126.

54. Yi H, Nevin KP, Kim BC, Franks AE, Klimes A, Tender LM, Lovley DR. 2009. Selection of a variant of *Geobacter sulfurreducens* with enhanced capacity for current production in microbial fuel cells. Biosens Bioelectron 24:3498–3503.

